# Development and Initial Characterization of Pigs with *DNAI1* Mutations and Primary Ciliary Dyskinesia

**DOI:** 10.1101/2024.05.22.594822

**Authors:** Mahmoud A. Abou Alaiwa, Brie M. Hilkin, Margaret P. Price, Nicholas D. Gansemer, Michael R. Rector, Mal R. Stroik, Linda S. Powers, Kristin M. Whitworth, Melissa S. Samuel, Akansha Jain, Lynda S. Ostedgaard, Sarah E. Ernst, Winter Philibert, Linda D. Boyken, Thomas O. Moninger, Phillip H. Karp, Douglas B. Hornick, Patrick L. Sinn, Anthony J. Fischer, Alejandro A. Pezzulo, Paul B. McCray, David K. Meyerholz, Joseph Zabner, Randy S. Prather, Michael J. Welsh, David A. Stoltz

## Abstract

Mutations in more than 50 different genes cause primary ciliary dyskinesia (PCD) by disrupting the activity of motile cilia that facilitate mucociliary transport (MCT). Knowledge of PCD has come from studies identifying disease-causing mutations, characterizing structural cilia abnormalities, finding genotype-phenotype relationships, and studying the cell biology of cilia. Despite these important findings, we still lack effective treatments and people with PCD have significant pulmonary impairment. As with many other diseases, a better understanding of pathogenic mechanisms may lead to effective treatments. To pursue disease mechanisms, we used CRISPR-Cas9 to develop a PCD pig with a disrupted *DNAI1* gene. PCD pig airway cilia lacked the outer dynein arm and had impaired beating. MCT was impaired under both baseline conditions and after cholinergic stimulation in PCD pigs. Neonatal PCD pigs developed neonatal respiratory distress with evidence of atelectasis, air trapping, and airway mucus obstruction. Despite airway mucus accumulation, lung bacterial counts were similar between neonatal wild-type and PCD pigs. Sinonasal disease was present in all neonatal PCD pigs. Older PCD pigs developed worsening airway mucus obstruction, inflammation, and bacterial infection. This pig model closely mimics the disease phenotype seen in people with PCD and can be used to better understand the pathophysiology of PCD airway disease.

## INTRODUCTION

Primary ciliary dyskinesia (PCD) is a genetic disease that causes airway mucus obstruction, inflammation, and infection of the pulmonary and sinonasal airways leading to worsening pulmonary function; sometimes necessitating lung transplantation (1–10). Mutations in more than 50 different genes cause PCD by disrupting the activity of motile cilia that facilitate mucociliary transport (MCT) (11, 12). Knowledge of PCD has come from studies identifying disease-causing mutations, characterizing structural cilia abnormalities, finding genotype-phenotype relationships, and studying the cell biology of cilia (1, 6, 11–23). Despite these findings, we still lack effective treatments, and people with PCD have significant upper and lower airway impairment (6, 19, 24).

As with nearly every other disease, a better understanding of pathogenic mechanisms can lead to effective treatments. Cystic fibrosis (CF) is an example (25). Using a CF pig model, that faithfully recapitulates human CF, we were able to investigate early disease timepoints not testable in humans (25–27). We discovered several key airway host defense defects that are present within hours of birth. In newborn CF pigs, loss of cystic fibrosis transmembrane conductance regulator (CFTR)-mediated chloride and bicarbonate transport led to: 1) a more acidic airway surface liquid (ASL) pH (28); 2) decreased airway antimicrobial factor killing due to the acidic ASL pH (28); 3) abnormal mucus viscoelastic properties (29); and 4) impaired MCT due to a failure of mucus strands to detach from submucosal gland (SMG) duct openings (30, 31). Additionally, we were able to answer the decades long chicken and egg conundrum about which comes first, inflammation or infection. We found that airway infection precedes airway inflammation in the CF lung (27, 32).

For PCD, we still lack answers to many key questions. For example: What is the underlying pathogenesis of neonatal respiratory distress in PCD? Does airway obstruction cause inflammation, in the absence of infection, or is inflammation preceded by infection? Is MCT abnormal at birth? What are the similarities and differences between upper (sinus) and lower airway disease in PCD? Two factors have limited progress in answering these questions. First, there has not been an animal model that fully replicates the typical sinonasal and lung disease that causes PCD morbidity and mortality. Second, there is an inability to study PCD pathogenesis at very early timepoints in life (e.g., the newborn/neonatal period). To overcome some of these limitations, we generated pigs with *DNAI1* mutations, hereafter, referred to as PCD pigs.

*DNAI1* mutations occur in approximately 10% of people with PCD and were the first genetic mutations identified to cause PCD in 1999 (33–35). *DNAI1* encodes a component of the outer dynein arm (ODA) of the central apparatus of motile cilia. A porcine model was chosen because the anatomy, physiology, and biochemistry of pig lungs are like those of humans (26). We were also encouraged to develop and study a porcine model of PCD based on the success of our earlier work with CF pigs (25). We report that PCD pigs have a significant impairment of cilia beating and decreased MCT. Shortly after birth, PCD pigs demonstrated features of neonatal respiratory distress and with time developed the hallmark features of the human disease including airway occlusion with mucus, inflammation, and infection.

## METHODS

### Generation of PCD pigs

To generate a porcine model of PCD, we targeted the porcine homolog of human *dynein axonemal intermediate chain 1* (*DNAI1*). *DNAI1* encodes a component of the outer dynein arm (ODA) of the central apparatus of motile cilia. The pig *DNAI1* gene, like human, has 20 exons (**Figure 1A**). There are mutation clusters in *DNAI1* exons 13, 16, and 17 (18, 34). The C-terminal half of the protein contains five WD repeat sequences (tryptophan and aspartic acid repeats) and the WD elements in DNAI1 are thought to mediate the assembly of the ODAs (36)}. The generation and identification of *DNAI1* KO piglets was carried out as described previously (37) with modifications specific to disrupting the *DNAI1* gene locus. We used CRISPR-Cas9 and 2 guide RNAs to introduce edits into pig exons 17 and 18, co-injecting 2 guide RNAs and Cas9 mRNA into porcine zygotes. The guide RNAs were designed to be injected as a pair to edit, and possibly remove, an area within the gene where a number of mutations in WD elements have been shown to result in defects in motile cilia function. The positions of guide RNAs are indicated as #1 and #2 in **Figure 1A**. Guide RNAs (sgRNAs) targeting exons 17 and 18 were identified using ChopChop v3 web tools (38): Guide 1, 5’-CACCTACGATGCCCACAACA-3’ (Exon 17); Guide 2, 5’-TATGACCTAAATTCGGCCGT-3’ (Exon 18). Guide RNAs were synthesized *in vitro* from Gene Block gene fragments (Integrated DNA Technologies, Coralville, IA) containing a T7 promoter sequence upstream of the 20 bp guide sequences (39). The guide RNAs, 10 ng/μl each, were mixed with s.p.Cas9 mRNA (TriLink BioTechnologies, San Diego, CA), 20 ng/μl final concentration, and injected into the cytoplasm of *in vitro* derived porcine zygotes generated from oocytes obtained from a Missouri abattoir. Embryos were cultured for 5 days (40–42) and morula- and/or blastocyst-stage embryos were transferred into a recipient sow.

**FIGURE 1.**
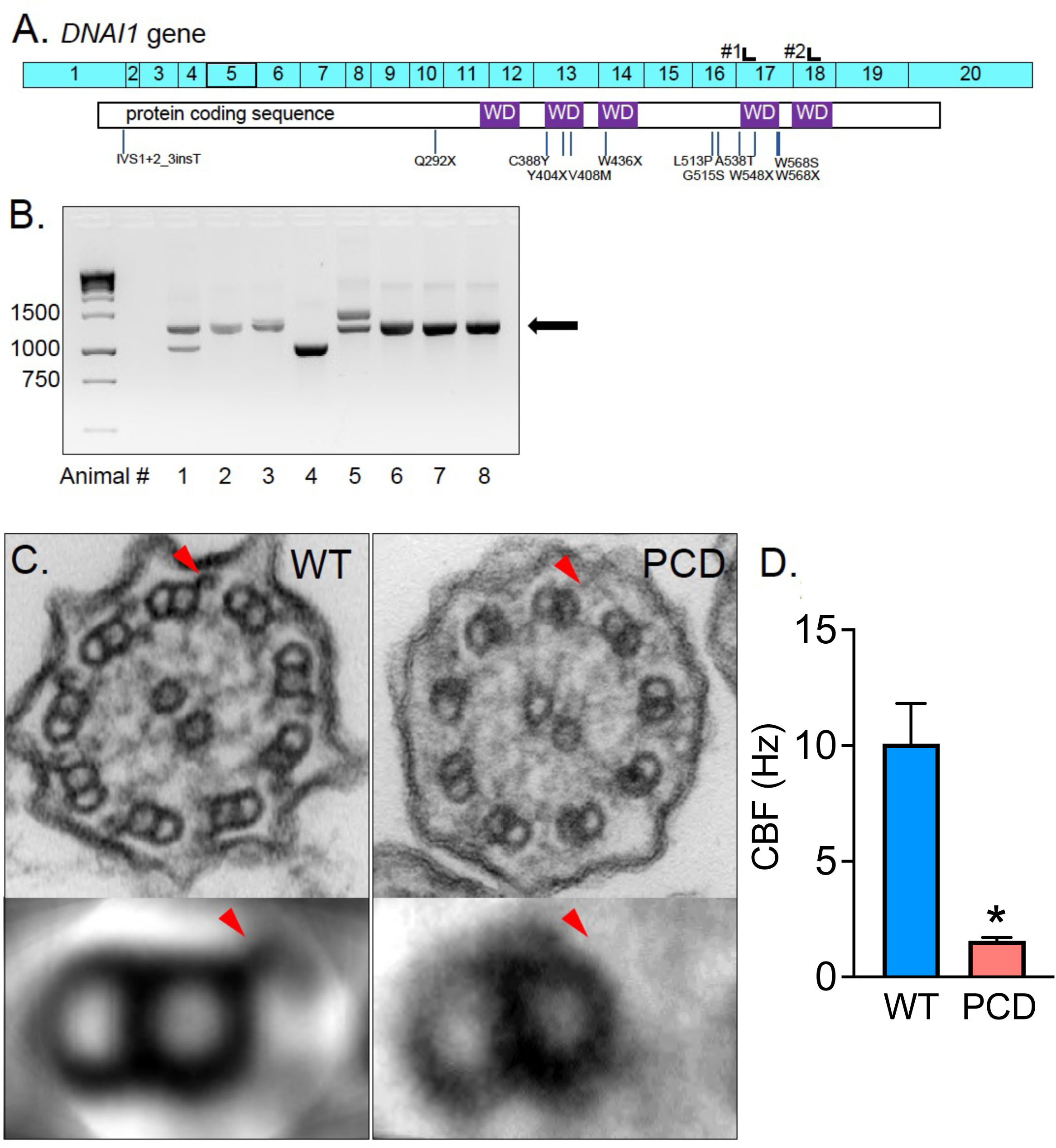
Generation of PCD Pigs by CRISPR-Cas9 Targeting of *DNAI1*. **(A)** Illustration of five conserved WD repeat regions relative to the 20 genomic exons mapped onto the encoded mRNA and protein coding sequence of the porcine *DNAI1* gene. IVS1+2_3insT is a recurring splice mutation and considered a common founder mutation. Locations of other mutations are shown. Position of guides #1 and #2 are shown. **(B)** PCR detection of the edited *DNAI1* gene in 8 piglets from a representative litter. Lane 7 shows the expected 1,230 WT bp band and lane 4 shows a biallelic deletion of the 226 bp between the guides to generate a 1,004 bp band. DNA sequencing revealed genomic mutations in the other pigs. Piglets 1-4 were female and 5-8 were male. **(C)** Image averaging of transmission emission tomography (TEM) of cilia from WT and PCD pigs. Arrowheads indicate outer dynein arm (ODA) position. Low and high power images, upper and lower panels, respectively. **(D)** Ciliary beat frequency (CBF) measurements in primary airway epithelial cultures from WT and PCD pigs. *n* = 8 WT and 15 PCD airway epithelial cultures from different animals. Error bars represent mean ± SEM. * *P* < 0.05 by Student’s *t*-test.

### Genotyping assay

Nucleotide primers flanking the expected CRISPR-Cas9 cut sites were designed using the Primer-Blast Program (43) and utilized to amplify the targeted fragments from lysates of piglet tail tissue. The tissue was lysed using the KAPA Express Extract Kit Plus Amplification Module and KAPA2G polymerase (KAPA Biosystems, Basel Switzerland). The forward primer (5’-GAT GTT TAT CAC TGT GGC AAG GAT TCA GT-3’) and reverse primer (5’-ATC GTT CTT TCT CAG CTT TCA TCA GTT CT-3’) amplify a 1,230 bp WT fragment.

PCR reaction conditions consisted of an initial denaturation at 95°C or 3 min, followed by 40 cycles of 95°C (15 s), 65°C (15 s), 72°C (30 s), and a final extension cycle of 72°C (1 min). The resulting PCR products were purified using the QIAquick PCR Purification Kit (Qiagen, Hilden, Germany), cloned into the TOPO TA vector, PCR 2.1-TOPO, and transfected into TOPO One Shot cells, and plated. Kanamycin-resistant colonies were picked and plasmids from ∼25 individual colonies were sequenced by Sanger sequencing to confirm the genotypes (Functional Biosciences, Madison, WI).

### Pig husbandry

After delivery, the piglets received colostrum. Piglets were either maintained on the sow or transitioned to milk replacer until mature enough to be placed on standard pig diets.

### TEM image acquisition and cilia ultrastructural analysis

Ciliary ultrastructure was evaluated on tracheal sections. Immediately following dissection, tissue was placed into fixative (2.5% glutaraldehyde in 0.1 M sodium cacodylate buffer, pH 7.2). After fixation, tissue was washed with 0.1 M phosphate buffer, stained with 1% osmium tetroxide, washed in H_2_O, stained with 2% uranyl acetate, washed in H_2_O, dehydrated through a graded series of ethanol and acetone and embedded in Spurr resin (EMS, Hatfield PA). Following a 24 hr polymerization at 60°C, 0.1 μm ultrathin sections were prepared and post-stained with lead citrate. Micrographs were acquired using a JEOL 1400+ transmission electron microscope (JEOL, Inc., Peabody, MA) operating at 80 kV equipped with a NanoSprint12 CMOS camera (AMT, Woburn, MA). PCD Detect was used as an image averaging technique to enhance visualization of ciliary ultrastructure features (44).

### Histopathological analyses

At necropsy, pigs underwent examination for gross lesions, and the observations were documented. Tissues were collected and preserved in 10% neutral buffered formalin, typically for 2-3 days. Subsequently, the tissues underwent routine processing, embedding, sectioning (approximately 4 μm thick), and staining with standard histochemical techniques such as hematoxylin and eosin (H&E) for general evaluation. Additionally, special stains were applied to specific tissues or lesions, including amylase-pretreated periodic acid-Schiff (dPAS) and modified Gram stain.

### Primary airway epithelial cell cultures

Epithelial cells were obtained from tracheas and nasal turbinates/ethmoids through enzymatic digestion. These cells were then seeded onto permeable filter supports and cultured at the air-liquid interface, following previously established protocols (45). The differentiated epithelia were utilized after a minimum of 14 days from seeding.

### BAL liquid collection and analyses

Bronchoalveolar lavage (BAL) was performed in neonatal and older pigs immediately after euthanasia and aseptic lung removal. An 8-ml aliquot of sterile saline was introduced into the mainstem bronchus, and this process was repeated twice. A comparable volume of liquid was retrieved from both WT and PCD neonatal piglets. BAL liquid was immediately placed on ice and transported to the laboratory for processing (cell counts and microbiology studies) and stored at −80°C for subsequent analysis. The total number of recovered cells in BAL liquid was quantified using a hemacytometer. Morphological differentiation of cells was carried out on cytospin preparations stained with the Differential Quik Stain Kit (Baxter, Deerfield, IL). Microbiologic studies were conducted on the collected BAL liquid.

### Microbiologic studies

Standard microbiologic techniques were utilized to quantify bacteria present in BAL liquid and lung tissue samples. The tissue samples were homogenized using a closed tissue grinder system. Both the homogenized tissue samples and the BAL liquid samples were serially diluted and plated onto Blood agar [tryptic soy agar with sheep blood (Remel, San Diego, CA)], Chocolate agar (Remel), and MacConkey agar (Remel). Organism identification was determined using a MALDI-TOF (matrix-assisted laser desorption/ionization time-of-flight) system (Bruker, Billerica, MA). Microbiology quantification data were logarithmically transformed. Prior to log transformation, 1 was added to all data values.

### Mucociliary transport studies

#### Ciliary beat frequency assay

Phase-contrast videos of ciliary motion were captured using a Zeiss Axio Observer microscope (50 frames per second) as previously described (46). For each field, we obtained 256 frames (5.1 seconds of video). We randomly selected a minimum of three fields per culture for measurement. To analyze the whole-field Gaussian mean ciliary beat frequencies (CBF), we utilized the Sisson-Ammons video analysis software (SAVA, Ammons Engineering, Mt. Morris, MI). Measurements were acquired at room temperature.

#### X-ray computed tomographic MCT assay

To measure MCT in the proximal airways, we used a previously published X-ray computed tomographic (CT) assay (31, 47). Pigs were sedated/anesthetized with ketamine (20 mg/kg, I.M., Dechra Veterinary Products) and acepromazine (0.2 mg/kg, I.M., Boehringer Ingelheim Animal Health USA, Inc.). IV dexmedetomidine (10 µg/kg/hr, I.V., Eugia US, LLC) was used to maintain anesthesia. The pigs were briefly intubated and tantalum microdisks (350 μm diameter x 25 μm thick, Sigma) were insufflated into the airways just beyond the vocal cords using a puff of air. Following delivery, the tubes and catheter were promptly removed. We obtained CT scans using a continuous spiral mode CT scanner (0.32 s rotation; 176 mm coverage in 1.5 s; 0.6 mm thick sections with 0.3 mm slice overlap, Siemens SOMATOM Force). Over a 6.3-min interval, we acquired forty-four CT scans. We tracked microdisk movement over time to obtain multiple measurements of their speed. We assessed microdisk clearance by determining whether a microdisk reached the larynx during the 6.3-min tracking period. The percentage of cleared microdisks was then calculated by dividing the number of cleared microdisks by the total number of microdisks tracked, expressed as a percentage.

### CT scan acquisition and analyses for lung and sinus disease

#### CT scan acquisition

Pigs underwent lung and sinus CT scanning using a scanning protocol like that previously described (48). Briefly, before scanning, pigs were anesthetized, intubated, mechanically ventilated, and then were paralyzed. After a lung recruitment maneuver, animals were scanned at 0 (expiratory scan) and 25 (inspiratory scan) cm H_2_O airway pressure. A Siemens Definition Flash CT scanner was used, and scanning parameters were as follows: 120 kVp, 40 mAs, slice thickness 0.6 mm, slice spacing 0.3 mm, and reconstruction kernel B60f. All CT scans were 512 × 512 voxels in the transverse plane. The newborn non-CF pig scans used for this study were retrospectively analyzed from scans acquired in an earlier study (49).

#### Lung analysis

Lung volume and airway measurements were quantified from CT DICOM data sets using the 3D Slicer software platform (http://www.slicer.org). Lung and airway volumes were obtained from automated segmentation. Endotracheal tube placement prohibited measurement of the more proximal trachea. Threshold values were used to categorize voxels as normal, air trapped (on expiratory CT scans), or atelectatic (on inspiratory CT scans) within the lung volume. Air trapped, normal, and atelectatic volumes were normalized to total lung volume and expressed as percentages.

#### Sinus analysis

CT DICOM datasets were imported into the 3D Slicer software platform (http://www.slicer.org) for sinus volume analysis, following a similar approach as previously described (50, 51). We utilized threshold values of voxels on the CT image to distinguish measurements of the skull and paranasal sinus volumes. Additionally, we manually segmented the ethmoid and maxillary sinuses. Specifically, the ethmoid sinus was defined as the sinus medial to the orbit, superior to the maxillary sinus, and posterior to the turbinates. The maxillary sinus was defined as the enclosed sinus lateral to the ethmoid, inferior to the orbit, and superior to the molars.

### Study approval

The University of Iowa and University of Missouri Institutional Animal Care and Use Committees approved all animal studies.

### Statistical analyses

Data are presented for individual animals with mean ± SEM. Differences were considered statistically significant at *P* < 0.05. Analyses were made in GraphPad Prism v10.2.3 (GraphPad Software, La Jolla, CA).

## RESULTS

### DNAI1 editing generated PCD pigs

Over 50 different cilia-related genetic mutations have been described to cause a PCD phenotype in humans. To generate PCD pigs, we targeted the porcine homolog of human *DNAI1* to produce loss of function mutations in pigs (hereafter referred to as PCD pigs). The pig *DNAI1* gene, like human, has 20 exons (**Figure 1A**). There are mutation clusters in human *DNAI1* exons 13, 16, and 17 (18, 34). The C-terminal half of the protein contains five WD repeat sequences. The WD elements in DNAI1 are thought to mediate the assembly of the ODAs (33, 36). The generation and identification of *DNAI1*-targeted piglets was carried out as described previously (37) with modifications specific to disrupting the *DNAI1* gene locus. We used CRISPR-Cas9 to edit an area within *DNAI1*where several mutations in WD elements have been shown to result in defects in motile cilia function. A pair of guide RNAs targeting exons 17 and 18 and *Cas9* mRNA were co-injected into the cytoplasm of *in vitro* derived porcine zygotes generated from oocytes (37).

Embryos were cultured for 5 days and morula- and/or blastocyst-stage embryos were transferred into a recipient sow (37). **Figure 1B** shows PCR detection of the edited *DNAI1* gene in 8 piglets from a representative litter. Lane 7 shows the expected 1,230 WT bp band and lane 4 shows a biallelic deletion of the 226 bp between the guides to generate a 1,004 bp band. DNA sequencing revealed genomic mutations in the other pigs. Piglets 1-4 were female and 5-8 were male.

### PCD pigs have impaired ciliary activity

Wild-type (WT) and PCD pigs had similar birth weights (1.2 ± 0.2 kg vs. 1.3 ± 0.1 kg) and there were a comparable number of male and female PCD pigs. Transmission electron microscopy (TEM) confirmed that PCD pigs lacked the ODA in motile cilia (**Figure 1C**). To test the hypothesis that ciliary function would be impaired in PCD pigs, we quantified ciliary beat frequency (CBF) in primary airway epithelial cell cultures grown at the air-liquid interface. CBF was significantly reduced in PCD cells compared to WT littermate controls (**Figure 1D**).

### PCD pigs have features of neonatal respiratory distress

Up to 80% of infants with PCD develop neonatal respiratory distress, typically within the first 12-48 hr following birth, due to lobar collapse/atelectasis which can require supplemental oxygen and prolonged hospitalization (10, 52–54). Therefore, we studied PCD piglets within the first days of birth to investigate for features of neonatal respiratory distress. Piglets studied within 24-36 hr of birth are defined as “36 hr” and those studied 36-72 hr are defined as “72 hr”.

We monitored pulse oximetry in a PCD pig litter immediately after birth. Half of the piglets developed transient hypoxemia in the first 72 hr. We performed high resolution computed tomography (CT) imaging of the lungs (inspiratory and expiratory scans) and sinuses of WT and PCD pigs to evaluate for atelectasis, air trapping, and sinus obstruction. Air trapping was significantly greater in PCD lungs, compared to control pigs (**Figure 2 and 3A**). Atelectasis was commonly observed in PCD pig lungs, with some pigs having regions of significant lung collapse (**Figure 2 and 3B**). In WT pigs, no atelectasis was observed. Neonatal PCD pig ethmoids, turbinates, and mastoids had areas of opacification, consistent with mucus accumulation (**Figure 4**). We quantified the maxillary and ethmoid sinus air volumes; both were significantly reduced in PCD pigs (**Figure 5**). These clinical and radiographic features are consistent with the neonatal respiratory distress observed in human newborns with PCD.

**FIGURE 2.**
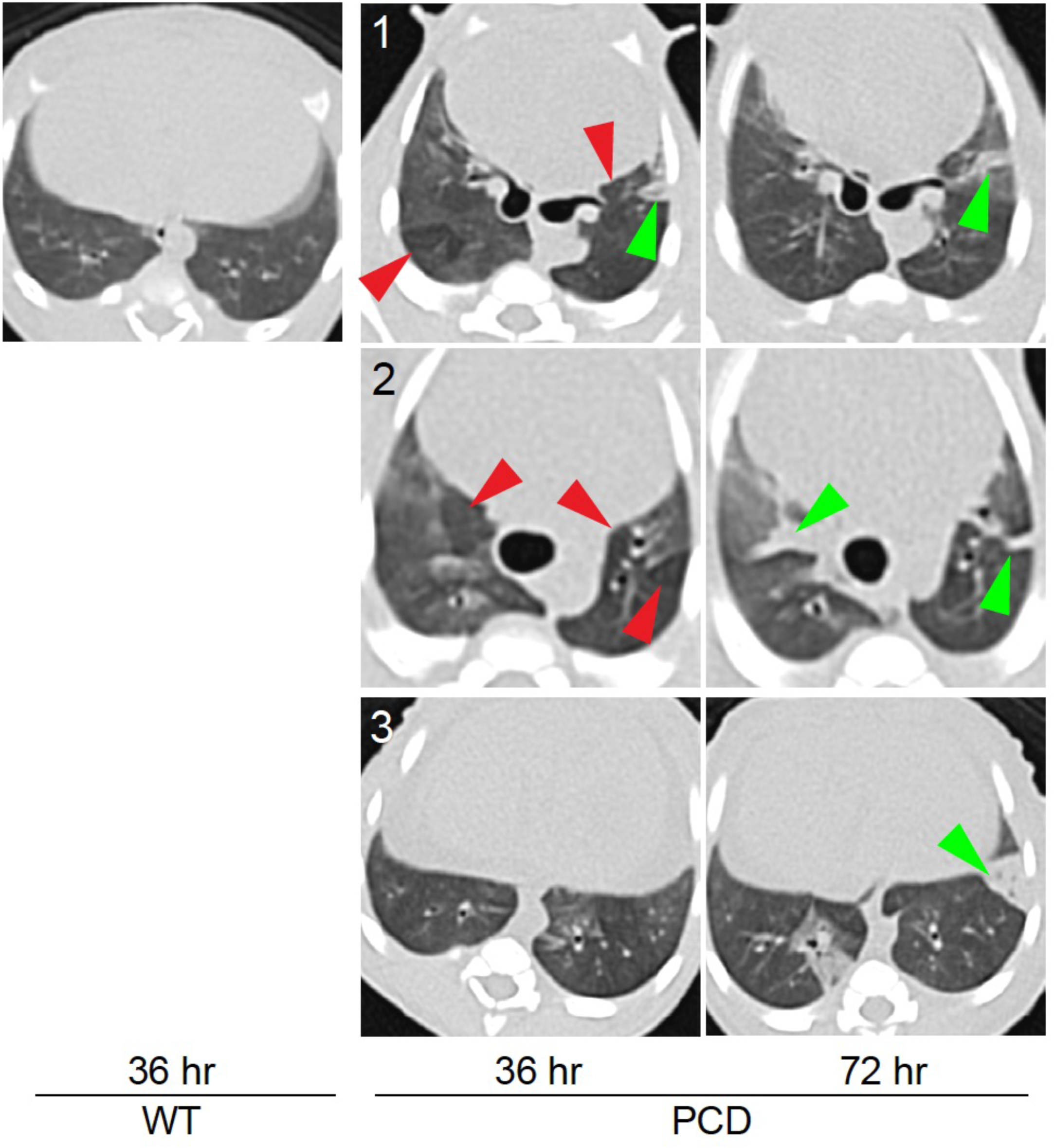
PCD Pigs Develop Radiographic Features of Neonatal Respiratory Distress Associated with Human PCD. CT images of the lungs acquired at ∼36 and 72 hr after birth from WT and PCD pigs. Representative images from 3 neonatal PCD pigs (#1-3) showing air trapping and atelectasis/lobar collapse. Red arrowheads denote regions of air trapping and green arrowheads denote regions of atelectasis/collapse.

**FIGURE 3.**
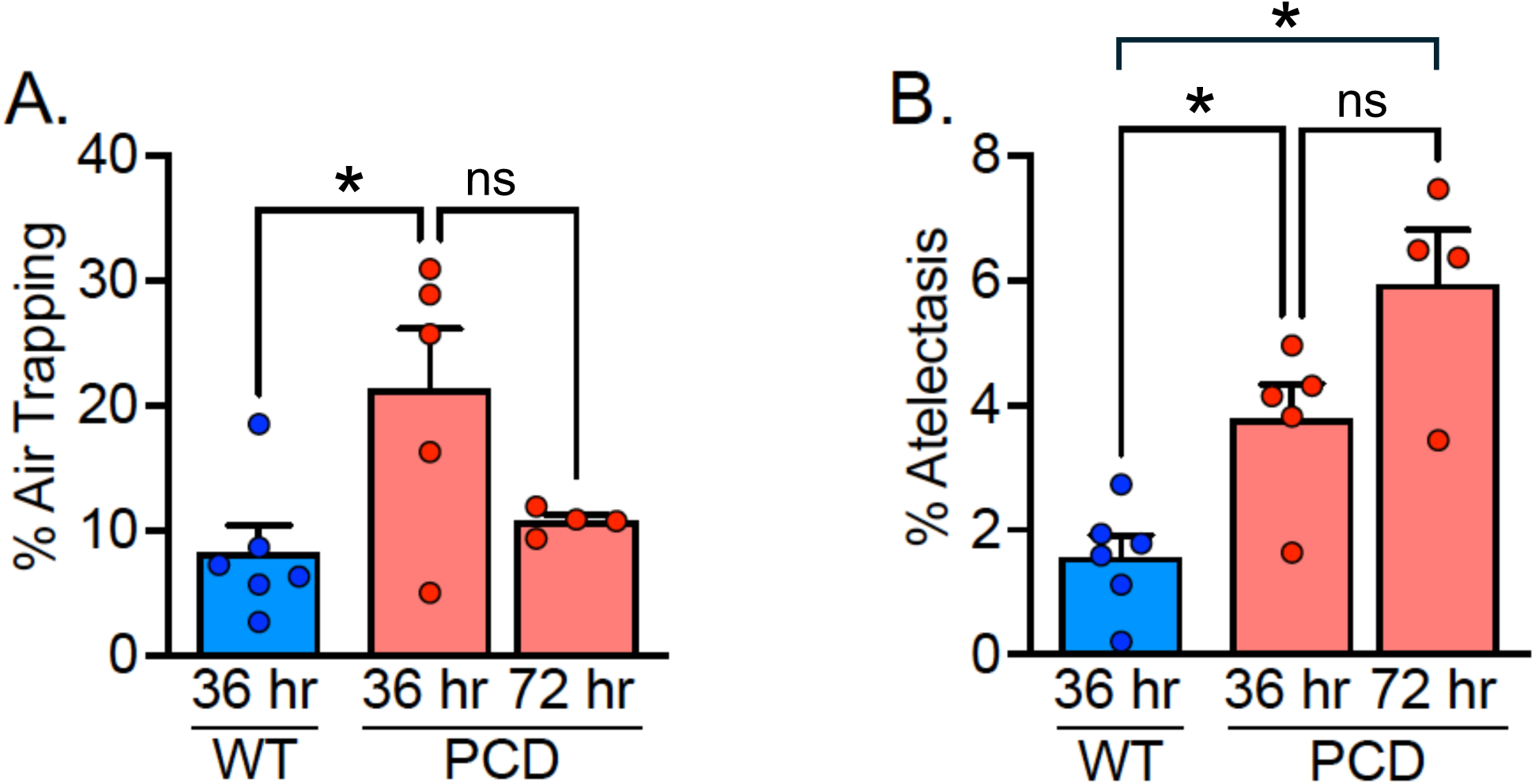
Air Trapping and Atelectasis are Present in PCD Pigs. Quantification of air trapping and atelectasis in WT and PCD pigs ∼36-72 hr old. Volumetric analyses were performed on the three-dimensional reconstructions of segmented chest CT scans. **(A)** Air trapping and **(B)** Atelectasis as a % of total lung voxels. Each symbol represents data from a different animal. Error bars represent mean ± SEM. * *P* < 0.05 by one-way ANOVA and Tukey’s multiple comparison test.

**FIGURE 4.**
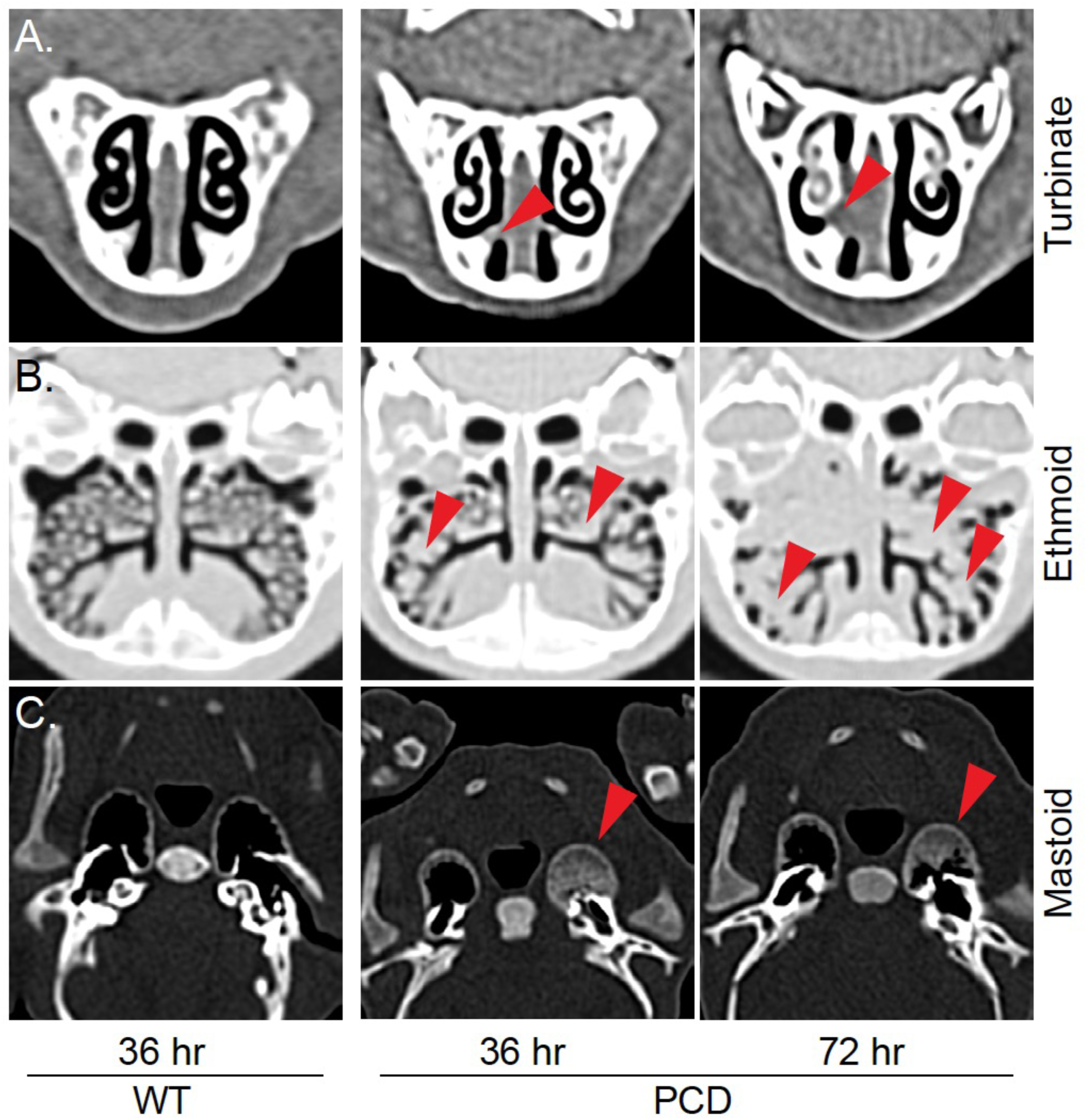
Sinus Obstruction Occurs Early in PCD Pigs. Data are coronal CT images of the upper airways acquired in WT and PCD pigs ∼36-72 hr old. **(A)** Turbinates and nasal passage. Arrowheads indicate mucosal thickening and mucus accumulation. **(B)** Ethmoid sinus. Arrowheads indicate mucus accumulation. **(C)** Mastoid air cells. Arrowheads indicate opacification of the air cells.

**FIGURE 5.**
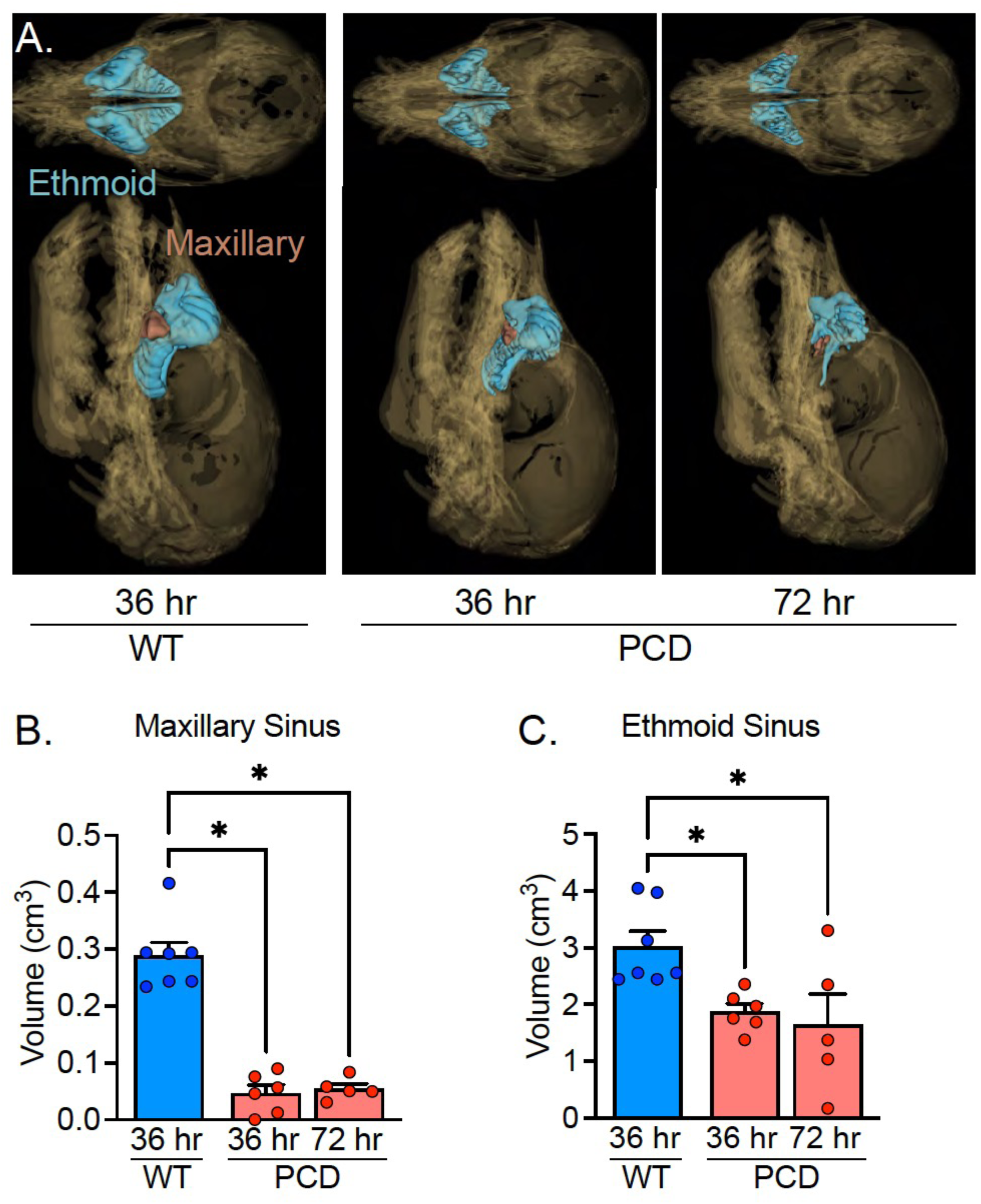
Sinus Air Volume is Reduced in Neonatal PCD Pigs. **(A)** Three-dimensional reconstruction view of the skull (transparent overlay) and sinuses with ethmoid (cyan) and maxillary (light red) sinus. **(B-C)** Volumetric analyses performed on the three-dimensional reconstructions of segmented sinus CT scans. Each symbol represents data from a different animal. Error bars represent mean ± SEM. * *P* < 0.05 by Kruskal-Wallis test with Dunn’s multiple comparisons test.

### Mucociliary transport is impaired in PCD pigs

Impaired ciliary motility suggested that mucociliary transport (MCT) would be disrupted in PCD pigs. To test this hypothesis, we measured MCT using an approach that we previously described (31, 47). We instilled tantalum microdisks (350 μm) into the proximal airways of WT and PCD neonatal piglets, performed high-resolution chest tomography (CT) scans (every 9 sec for 6.3 min), and measured microdisk clearance from the lungs. Studies were performed under both control conditions and following methacholine stimulation, to increase ciliary beating and submucosal gland secretion. In WT pigs, baseline microdisk clearance was approximately 25% and following cholinergic stimulation increased to ∼75% (**Figure 6**). PCD pigs failed to clear any microdisks under baseline conditions or after cholinergic stimulation during the scanning period (**Figure 6**).

**FIGURE 6.**
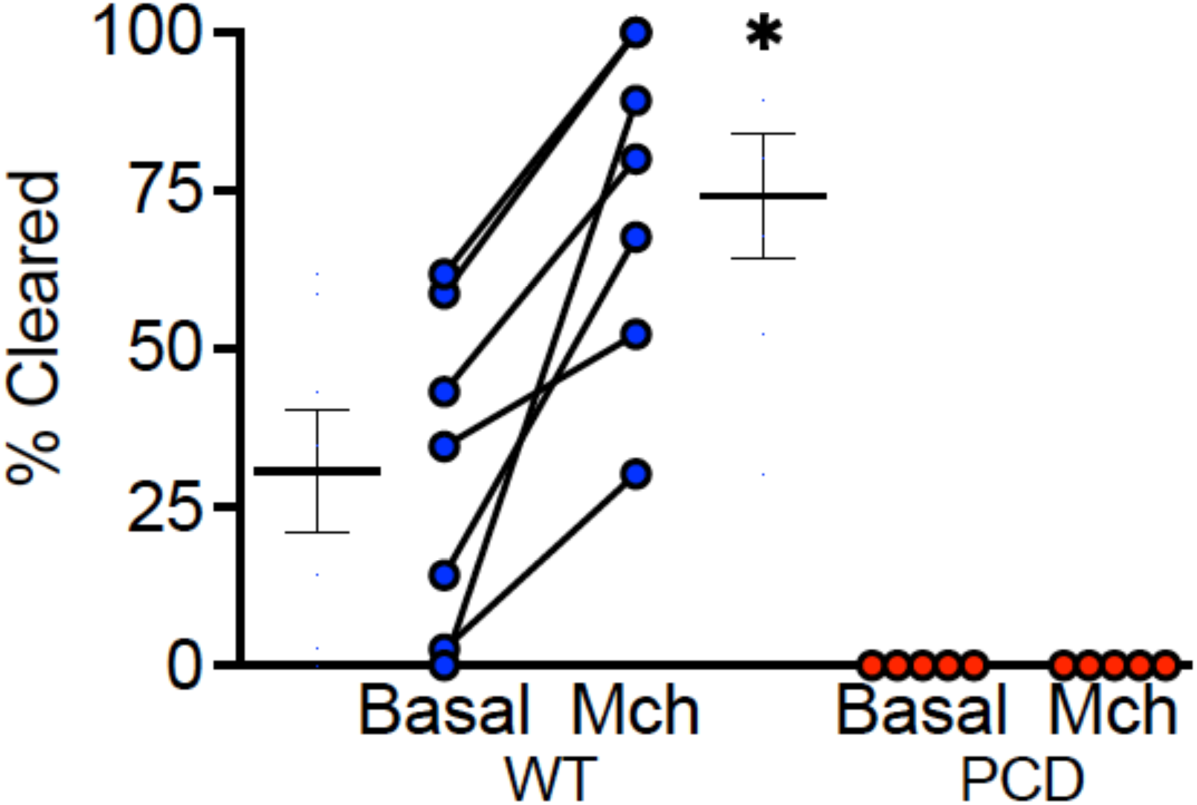
Mucociliary Transport is Significantly Reduced in PCD Pigs. Microdisk clearance was measured with a CT-based assay. Clearance was measured under baseline conditions and after methacholine stimulation. Each symbol represents data from a different animal. Error bars represent mean ± SEM. * *P* < 0.05 by paired Student’s *t*-test.

### Neonatal PCD pig lungs and sinuses have mucus obstruction and inflammation, but lack significant infection

To assess for airway disease in PCD pigs, we necropsied a cohort of neonatal pigs after CT scans were performed. We found no evidence of *situs inversus* in PCD pigs. At necropsy, PCD pig sinuses were nearly always filled with mucus (**Figure 7**). In the lungs, the airways were variably obstructed with mucus, and had additional cellular debris present with oftentimes adjacent lung regions that appeared collapsed (**Figure 8**). Similar findings were observed in tissue from the sinuses (**Figure 7**).

**FIGURE 7.**
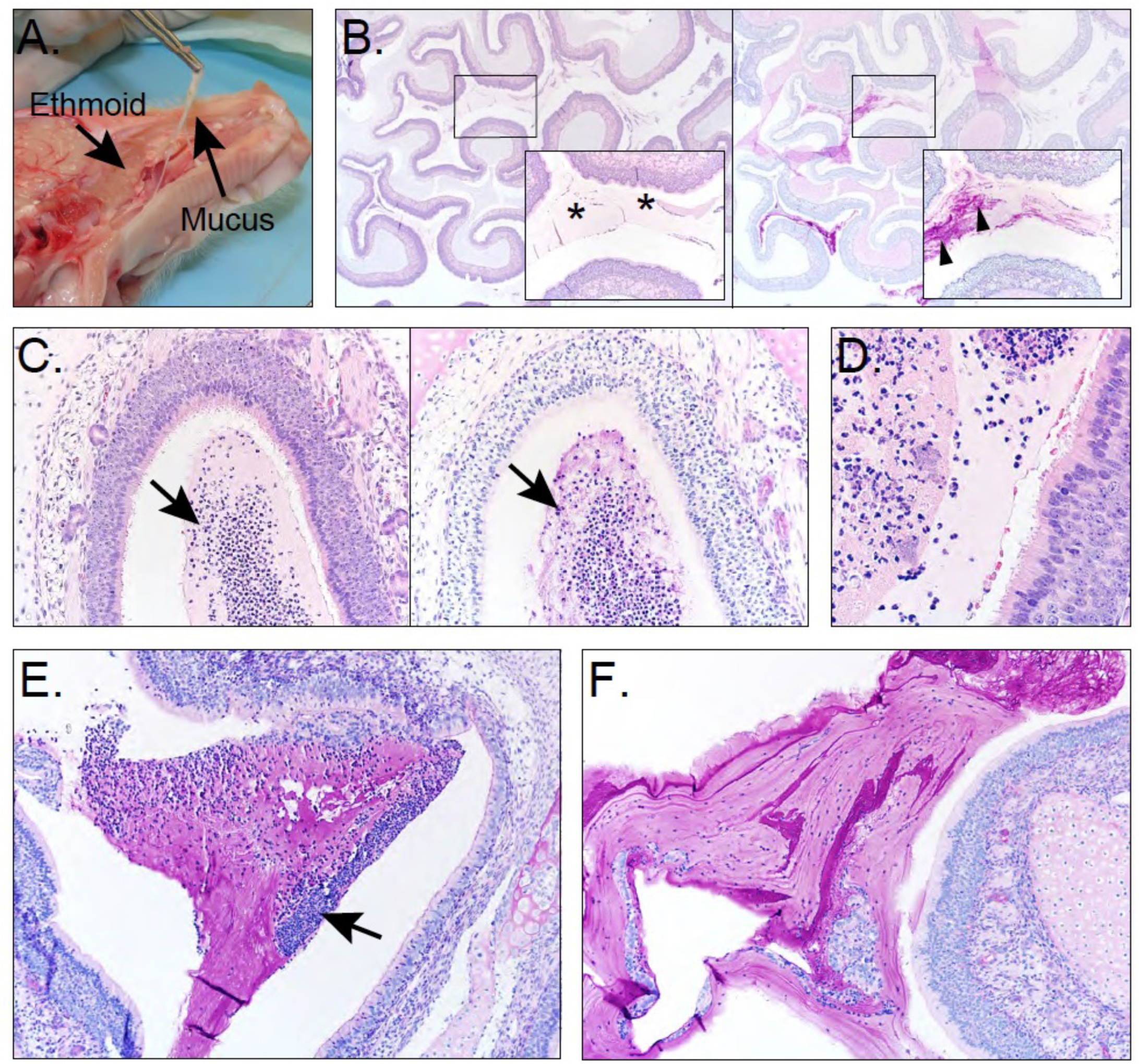
Neonatal PCD Pig Sinuses Have Mucus Accumulation and Inflammation. **(A)** Gross image of tenacious mucus pulled out from the crevices of the ethmoid sinus in a ∼24 hr old PCD pig. **(B-C)** Microscopic image of the ethmoid sinus, H&E (left) and dPAS (right). (B) Eosinophilic secretions in lumen of H&E sections (star, left inset) were partially dPAS positive (mucus, arrowhead, right inset). 2x and 10x (inset) magnification. (C) Sinusitis with mucopurulent exudate (arrows). 2x magnification. **(D)** Mucopurulent sinusitis of the ethmoid. H&E sections, 40x magnification. **(E-F)** dPAS microscopic sections of the ethmoid sinus at 10x magnification. (E) Mucopurulent focus with severe neutrophilic infiltration (arrow). (F) Mucus with scattered minimal neutrophils.

**FIGURE 8.**
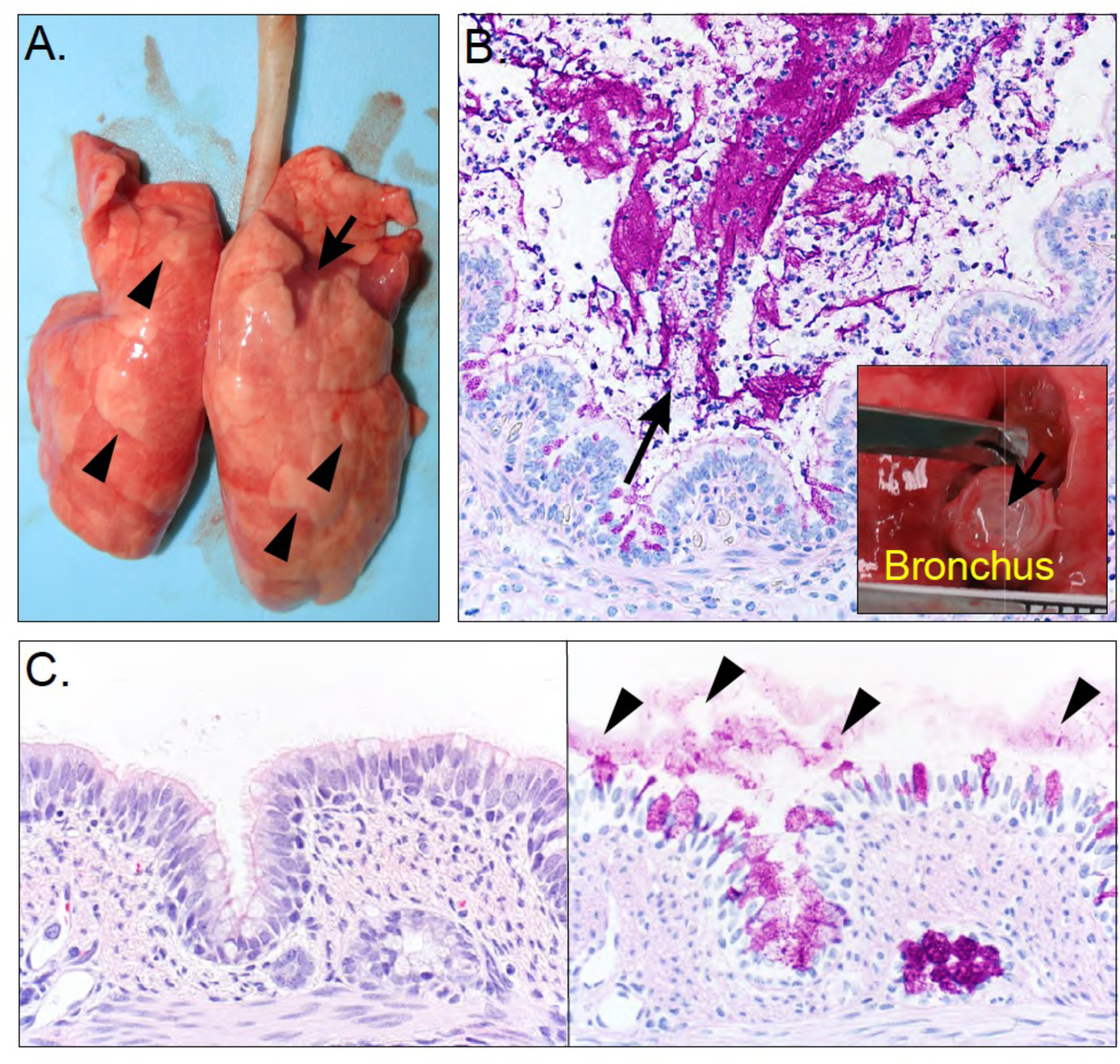
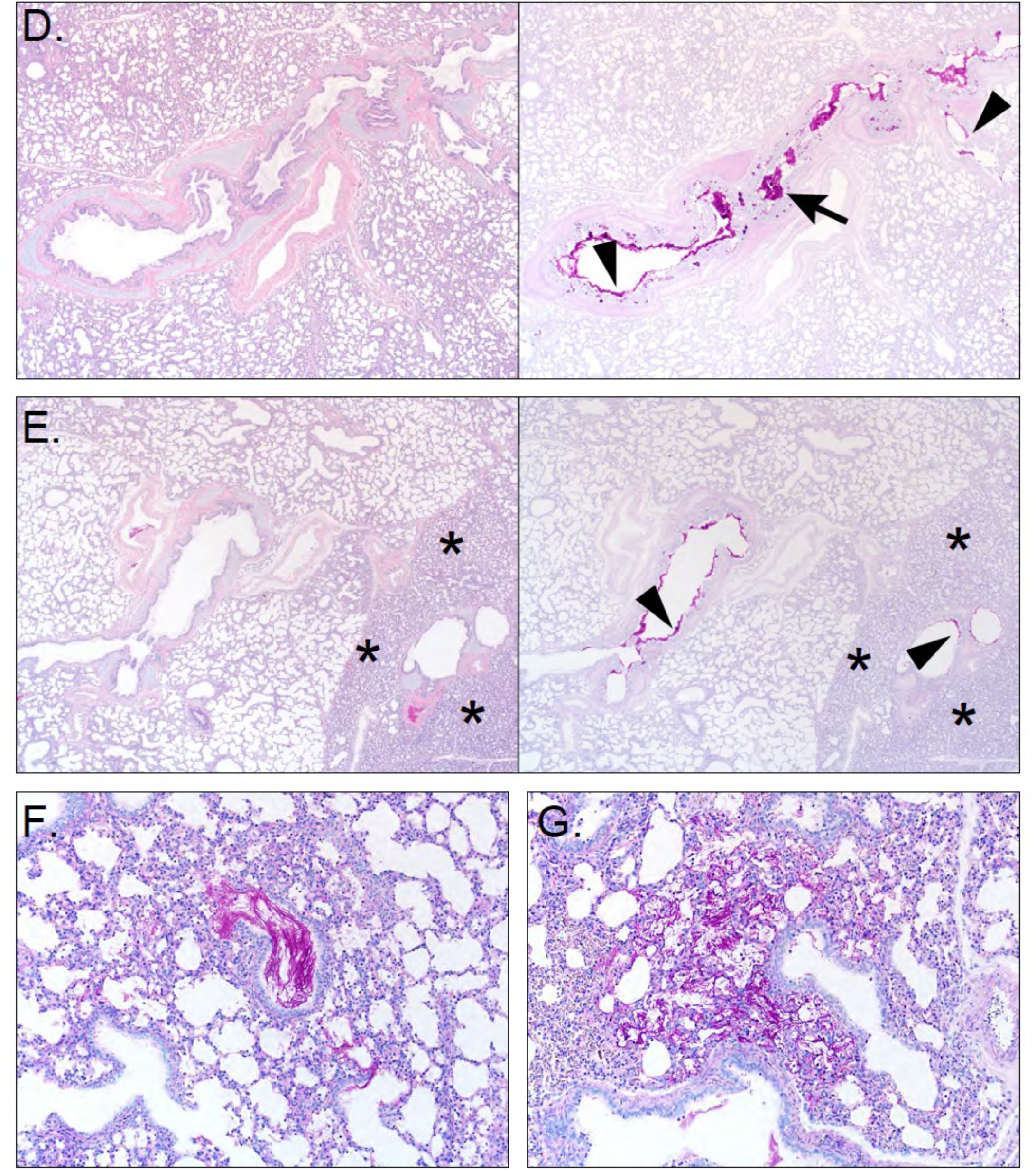
Mucus Accumulation Characterizes the Lung Abnormalities in PCD Pigs. **(A)** Gross image of air trapping (arrowheads) and lobar collapse (arrow) in the lungs of a ∼24 hr old PCD pig. **(B)** dPAS microscopic section of a large airway at 20x magnification with mucus, neutrophils and cellular debris accumulated in the lumen (arrow). Inset is a gross image of a bronchus section showing mucus plugging the lumen (arrow). **(C)** Microscopic image of the bronchus at 40x magnification, H&E (left) and dPAS (right), showing a rim of mucoid material with no inflammation (arrowheads). **(D-E)** Microscopic image of the airways at 2x magnification, H&E (left) and dPAS (right). (D) Bronchus, dPAS+ mucus (arrowheads) can be seen as a rim along the surface epithelia to overt obstruction of airway (arrow). (E) Small airway, dPAS+ mucus (arrowheads) can be seen covering the surface epithelium with atelectasis (asterisks). **(F-G)** dPAS microscopic sections of the lungs at 10x magnification. (F) Multifocal mucus in scattered large, small airways, and alveoli. (G) Alveoli filled up with mucus.

During the neonatal period, CF pigs already have more bacteria in their lungs than WT pigs (32). We expected to observe a similar pattern in neonatal PCD pigs. Instead, we found comparable numbers of bacteria in the lungs of neonatal WT and PCD pigs (**Figure 9A,B**). This was the case irrespective of sampling method: lung tissue homogenate versus bronchoalveolar lavage (BAL) fluid. Similar bacterial species were recovered from WT and PCD pig lungs (**Figure 9C,D**). In summary, neonatal PCD pigs had evidence of both upper and lower airway disease with mucus plugging. This phenotype is consistent with neonatal respiratory distress, seen in infants with PCD, and suggest that the underlying etiology is airway mucus obstruction. Despite these changes, we did not observe more bacteria in neonatal PCD pigs.

**FIGURE 9.**
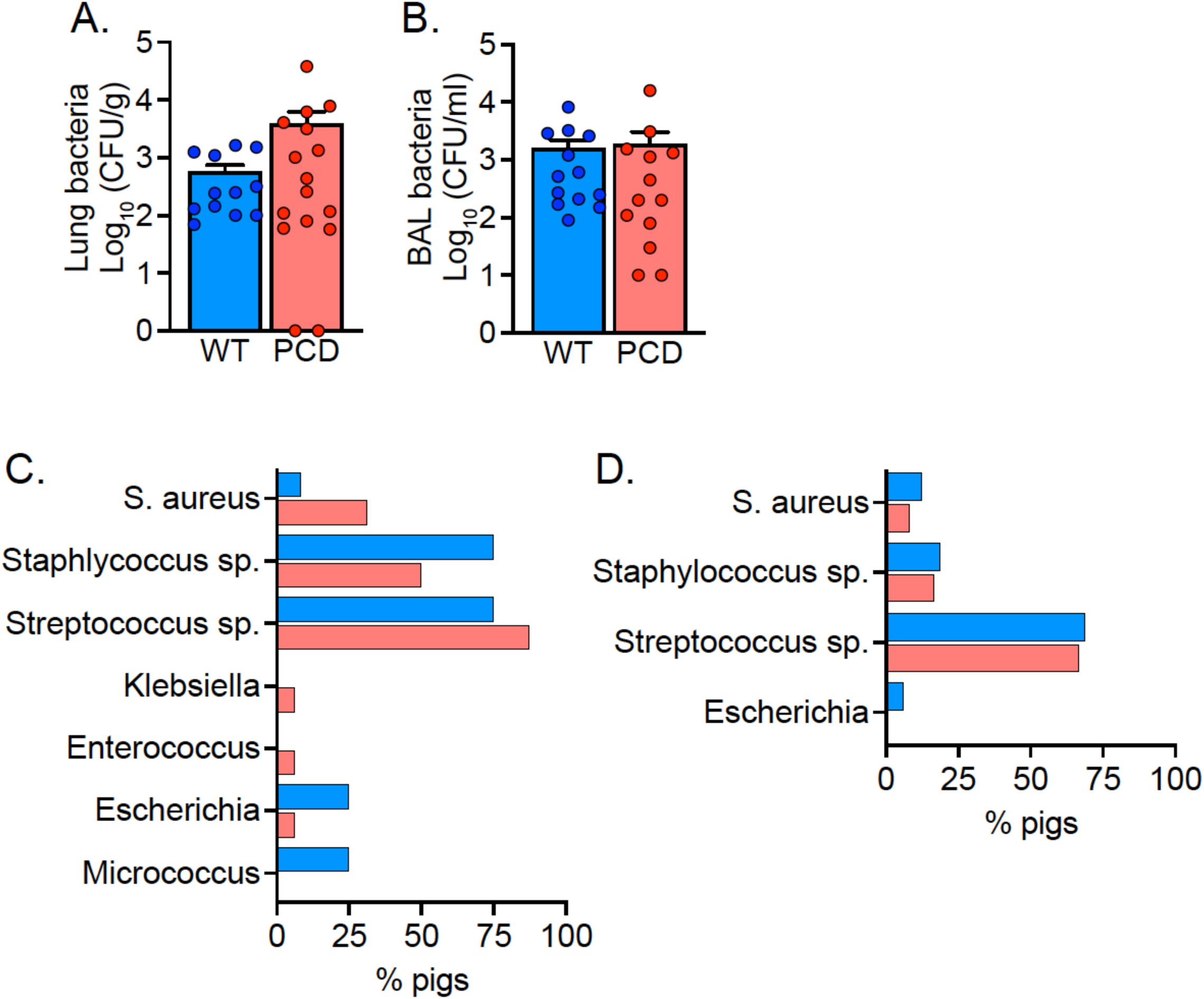
Neonatal PCD Pig Lungs are Not Infected. **(A)** Lung bacterial counts. **(B)** Bronchoalveolar lavage (BAL) liquid bacterial counts. **(C-D)** Percentage of bacterial species in (C) lung homogenates and (D) BAL liquid. WT (blue bars) and PCD (red bars) pigs were all less than 72 hr old. Each symbol represents data from a different animal. Error bars represent mean ± SEM.

### PCD pigs develop progressive lung and sinus disease over time

To determine if older PCD pigs develop disease like humans, we studied 6 to 12-month-old PCD pigs. Mucus obstruction and inflammation were present in the sinuses (**Figure 10A-C**). On gross examination, the lungs had areas of atelectasis/collapse with mucus obstructed airways (**Figure 11A-C**). Lung histology revealed changes of chronic inflammation and bronchiectasis (**Figure 11D-F**). We found increased neutrophils in PCD BAL liquid (**Figure 12A**) and greater bacteria in PCD lungs (**Figure 12B,C**). The isolated bacteria included *Pseudomonas aeruginosa*, *Trueperella pyogenes*, *Rothia nasimurium*, and *Corynebacterium xerosis*. These pulmonary findings replicate those in older PCD patients.

**FIGURE 10.**
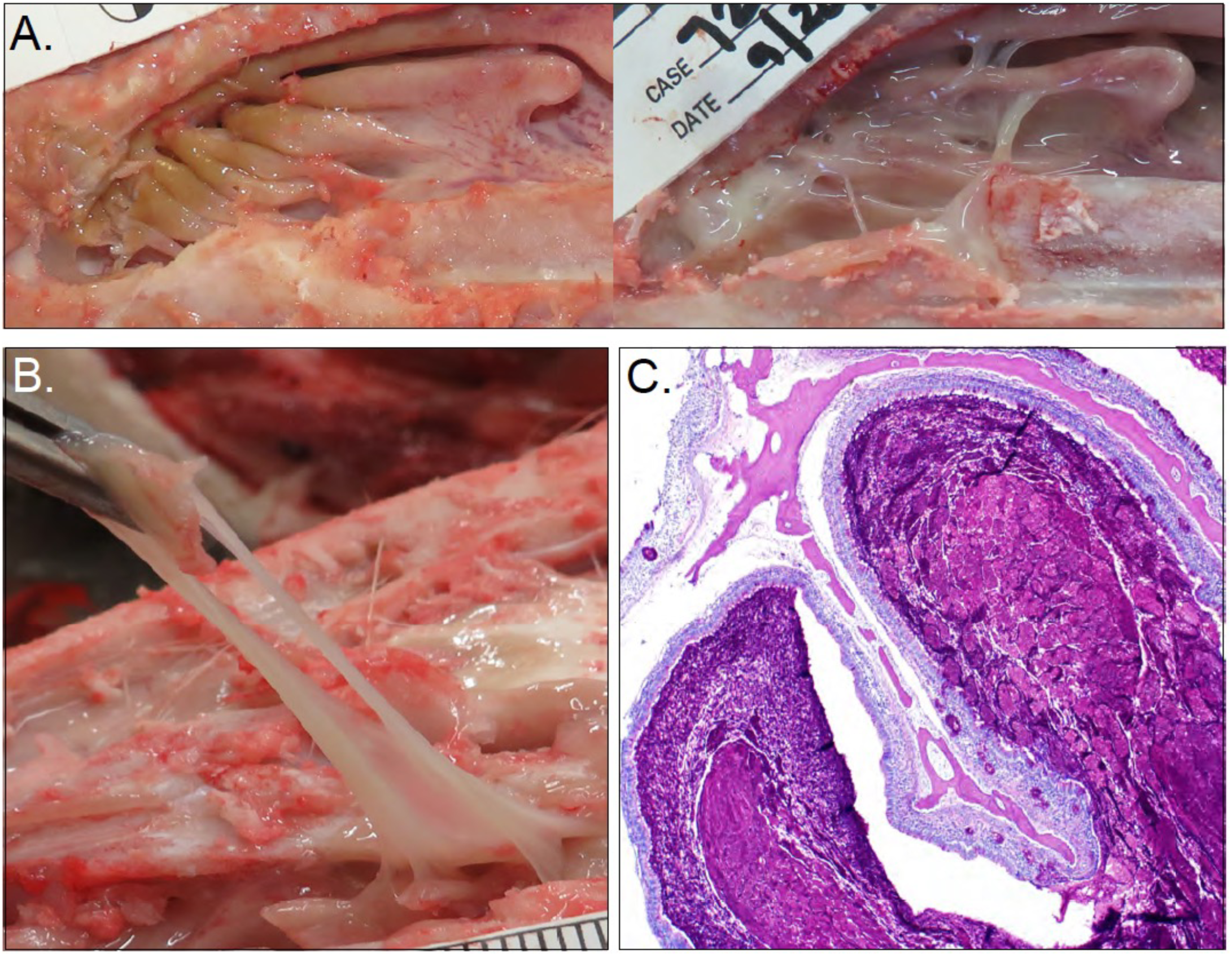
PCD Pigs Develop Progressive Sinus Disease. **(A)** Gross images of ethmoid sinuses. WT – left panel. PCD – right panel. **(B)** Thick mucus in the PCD pig ethmoid sinus. **(C)** dPAS image showing mucopurulent sinus disease in PCD pigs. 1 yr-old PCD pig.

**FIGURE 11.**
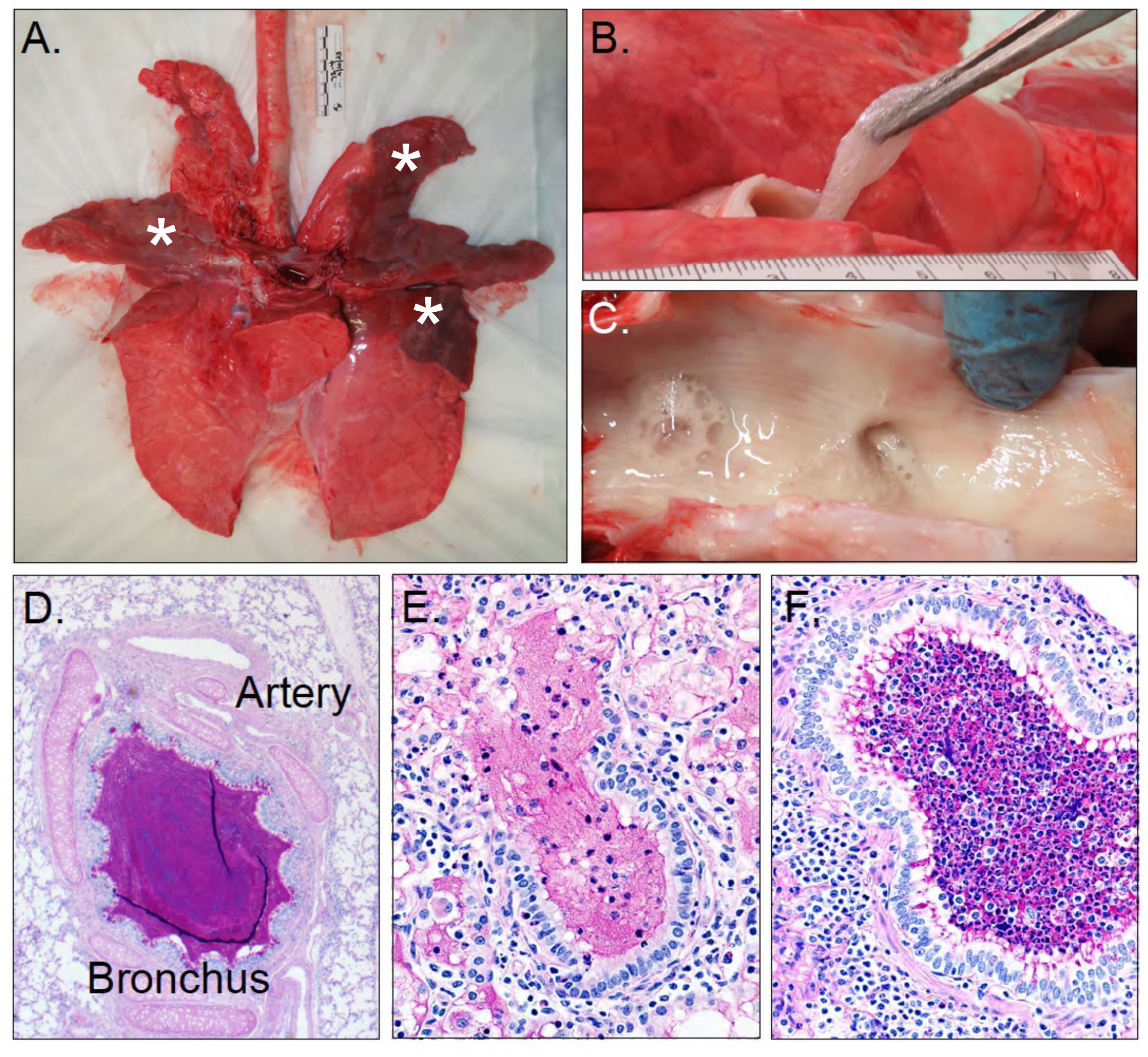
Older PCD Pigs Develop Airway Disease Similar to Human PCD. **(A-C)** Gross images of the lungs of a 1 yr-old PCD pig. (A) Asterisks denote areas of lung collapse/atelectasis. (B) Mucus in trachea. (C) Mucus emerging from tracheal bronchus. **(D)** A dilated/bronchiectatic airway filled with dPAS material, 6 month-old PCD pig. **(E-F)** dPAS filled airway lumens with variable levels of neutrophilic infiltration, 9 month-old PCD pig.

**FIGURE 12.**
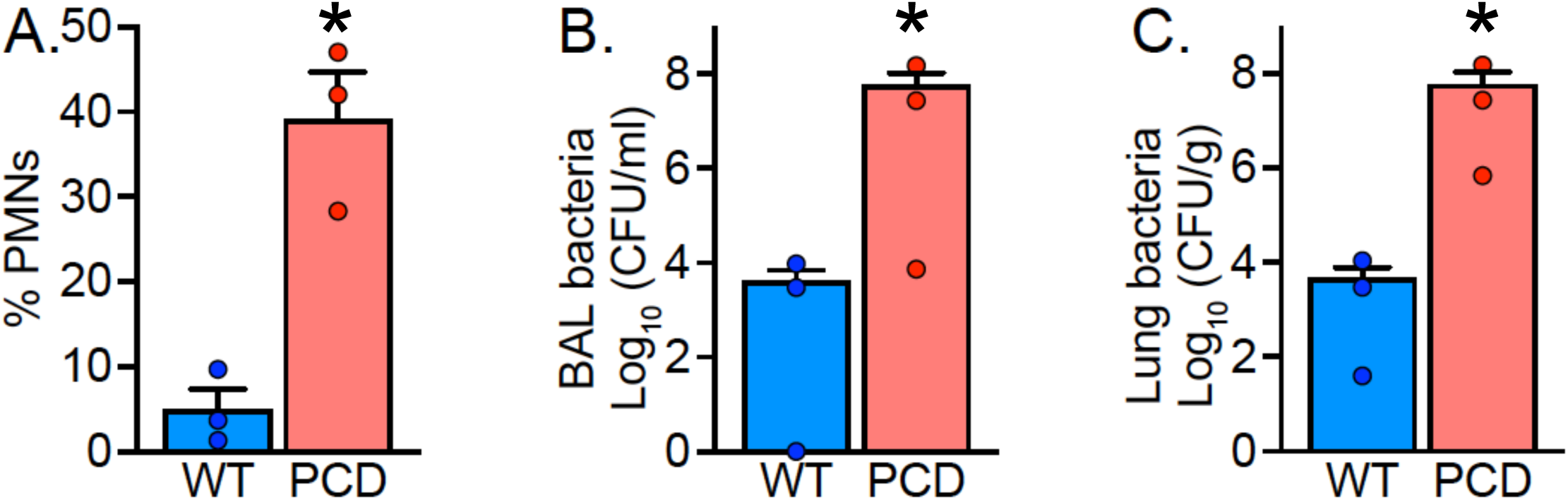
Older PCD Pigs Develop Airway Inflammation and Infection. (A) **BAL liquid % PMNs.** (B) **BAL liquid bacterial counts.** (C) Lung homogenate bacterial counts. Each symbol represents data from a different animal. WT and PCD pigs were studied at 6, 9, and 12 months old. Error bars represent mean ± SEM. * *P* < 0.05.

## DISCUSSION

We disrupted *DNAI1* with the CRISPR-Cas9 system to generate a porcine model of PCD. PCD pigs exhibited hallmark features of human PCD sinus and lung disease, including neonatal respiratory distress, a clinical feature observed in up to 80% of infants with PCD (10, 53–55). In neonatal PCD pigs, the sinuses were obstructed and atelectasis as well as air trapping were present in the lungs. These changes were likely due, in part, to mucus accumulation in the upper and lower airways. With time, airway disease progressed in PCD pigs that included airway infection and bronchiectasis.

We had the unique opportunity to investigate the pathogenesis of PCD at early timepoints not previously feasible in infants with PCD or murine PCD models. Our findings demonstrate that the consequences of disrupted ciliary function are apparent shortly after birth. By studying newborn PCD pigs, we discovered the following about the early pathogenesis of PCD. First, in infants with PCD, the etiology of neonatal respiratory distress has been postulated to be impaired fetal lung liquid clearance leading to atelectasis/lobar collapse (56, 57). However, this assumption has not been directly tested. In contrast to transient tachypnea of the newborn, which occurs immediately after birth and likely is related to impaired lung liquid clearance, neonatal respiratory distress tends to occur 12 hours or later after birth (54). Our data suggest that the primary mechanism of PCD-related neonatal respiratory distress is mucus obstruction in the airways, and less likely related to defects in lung liquid clearance. Second, newborn PCD pigs had significant airway mucus obstruction, yet lacked bacterial infection. Third, proximal airway MCT was absent in newborn PCD pigs. Fourth, airway obstruction seemed to be more severe in the sinuses of neonatal PCD pigs, than in the lungs, suggesting that other clearance mechanisms in the lung, like cough clearance, remain intact.

With time, PCD and CF have similar clinical features including chronic airway infection, inflammation, and mucus obstruction; sinusitis; bronchiectasis; and a progressive decline in pulmonary function. In contrast to PCD, newborns with CF rarely present with respiratory abnormalities during the neonatal period. Our PCD and CF pigs mirror these clinical features. By studying both PCD and CF pigs, we can begin to better understand the similarities and differences in pathogenesis between these two diseases. The current study already suggests several conclusions. First, CF is not solely due to impaired MCT. If it were, then the disease phenotype between CF and PCD should be similar. CF has multiple airway host defense defects, while PCD is primarily due to defective MCT. We previously discovered that, within hours of birth, CF pigs had more bacteria in their lungs compared to non-CF (32). Thus, we were surprised to find a similar number of bacteria between neonatal WT and PCD pigs. These findings suggest that, in the PCD airways, other host defense defects, such as antimicrobial factor killing, remain intact; at least at very early timepoints in the disease. Additionally, cough clearance also likely remains a very important airway host defense mechanism. Second, the MCT defect in PCD is more severe than in CF. In newborn CF pigs, under baseline conditions, MCT is normal (31). However, after cholinergic stimulation, mucus strands fail to detach from CF submucosal gland ducts thus impairing MCT (31). In contrast, MCT was impaired both under baseline conditions and after cholinergic stimulation in newborn PCD pigs. Third, sinus disease appears to be more severe in PCD than CF. This could be explained by the fact that the upper airways/sinuses lack other mechanisms to clear mucus including cough and two-phase gas-liquid flow (58–60).

Our study has strengths and limitations. Strengths include: 1) Porcine airway anatomy and physiology is like humans. 2) PCD pigs mimic the hallmark features of human PCD airway disease. 3) We studied PCD pigs shortly after birth, timepoints previously not testable. 4) We were able to compare concurrently MCT, CT imaging, bronchoalveolar lavage samples, microbiology, and histopathology. 5) PCD pigs develop both upper and lower airway disease. 6) *In vivo* and *in vitro* studies were possible using tissue from the same animals. 7) We studied pigs with a common human PCD-related mutation. 8) PCD pigs did not develop hydrocephalus which would have limited the usefulness of this model.

Limitations include: 1) The potential role of respiratory viruses in disease pathogenesis was not addressed in this study. However, our pigs have limited exposure to respiratory viruses under their current housing conditions. 2) The PCD pig’s response to a bacterial challenge was not investigated. This will be a focus of future studies. 3) We did not determine whether otitis media was present in PCD pigs. However, mastoiditis was a common feature. 4) We only used culture-dependent methods for microbiology studies. Whether culture-independent methods would be useful for further understanding disease pathogenesis is unknown.

In summary, PCD pigs provide a unique opportunity to better understand early disease pathogenesis which could lead to novel therapeutics, preventive treatments, and/or clinical trial endpoints/assays.

## ACKNOWLEDGEMENTS

This work was supported, in part, by NIH (T32 HL007638 and P30 DK054759) and the University of Iowa Distinguished Scholars Program.

## REFERENCES

1. Horani A, and Ferkol TW. Understanding Primary Ciliary Dyskinesia and Other Ciliopathies. J Pediatr. 2021;230:15–22 e1.

2. Shapiro AJ, Zariwala MA, Ferkol T, Davis SD, Sagel SD, Dell SD, et al. Diagnosis, monitoring, and treatment of primary ciliary dyskinesia: PCD foundation consensus recommendations based on state of the art review. Pediatr Pulmonol. 2016;51(2):115–32.

3. Leigh MW, Horani A, Kinghorn B, O’Connor MG, Zariwala MA, and Knowles MR. Primary Ciliary Dyskinesia (PCD): A genetic disorder of motile cilia. Transl Sci Rare Dis. 2019;4(1-2):51–75.

4. Shapiro AJ, Davis SD, Polineni D, Manion M, Rosenfeld M, Dell SD, et al. Diagnosis of Primary Ciliary Dyskinesia. An Official American Thoracic Society Clinical Practice Guideline. Am J Respir Crit Care Med. 2018;197(12):e24–e39.

5. Fahy JV, and Dickey BF. Airway mucus function and dysfunction. N Engl J Med. 2010;363(23):2233–47.

6. Leigh MW, Pittman JE, Carson JL, Ferkol TW, Dell SD, Davis SD, et al. Clinical and genetic aspects of primary ciliary dyskinesia/Kartagener syndrome. Genet Med. 2009;11(7):473–87.

7. Leigh MW, Ferkol TW, Davis SD, Lee HS, Rosenfeld M, Dell SD, et al. Clinical Features and Associated Likelihood of Primary Ciliary Dyskinesia in Children and Adolescents. Ann Am Thorac Soc. 2016;13(8):1305–13.

8. Date H, Yamashita M, Nagahiro I, Aoe M, Andou A, and Shimizu N. Living-donor lobar lung transplantation for primary ciliary dyskinesia. Ann Thorac Surg. 2001;71(6):2008–9.

9. Marro M, Leiva-Juarez MM, D’Ovidio F, Chan J, Van Raemdonck D, Ceulemans LJ, et al. Lung Transplantation for Primary Ciliary Dyskinesia and Kartagener Syndrome: A Multicenter Study. Transpl Int. 2023;36:10819.

10. Wee WB, Kinghorn B, Davis SD, Ferkol TW, and Shapiro AJ. Primary Ciliary Dyskinesia. Pediatrics. 2024.

11. Brennan SK, Ferkol TW, and Davis SD. Emerging Genotype-Phenotype Relationships in Primary Ciliary Dyskinesia. Int J Mol Sci. 2021;22(15).

12. Kinghorn B, Rosenfeld M, Sullivan E, Onchiri F, Ferkol TW, Sagel SD, et al. Airway Disease in Children with Primary Ciliary Dyskinesia: Impact of Ciliary Ultrastructure Defect and Genotype. Ann Am Thorac Soc. 2023;20(4):539–47.

13. Mitchison HM, and Smedley D. Primary ciliary dyskinesia: a big data genomics approach. Lancet Respir Med. 2022;10(5):423–5.

14. Lucas JS, Davis SD, Omran H, and Shoemark A. Primary ciliary dyskinesia in the genomics age. Lancet Respir Med. 2020;8(2):202–16.

15. Niziolek M, Bicka M, Osinka A, Samsel Z, Sekretarska J, Poprzeczko M, et al. PCD Genes-From Patients to Model Organisms and Back to Humans. Int J Mol Sci. 2022;23(3).

16. Lee L, and Ostrowski LE. Motile cilia genetics and cell biology: big results from little mice. Cell Mol Life Sci. 2021;78(3):769–97.

17. Solomon GM, Francis R, Chu KK, Birket SE, Gabriel G, Trombley JE, et al. Assessment of ciliary phenotype in primary ciliary dyskinesia by micro-optical coherence tomography. JCI Insight. 2017;2(5):e91702.

18. Zariwala MA, Knowles MR, and Omran H. Genetic defects in ciliary structure and function. Annu Rev Physiol. 2007;69:423–50.

19. Davis SD, Rosenfeld M, Lee HS, Ferkol TW, Sagel SD, Dell SD, et al. Primary Ciliary Dyskinesia: Longitudinal Study of Lung Disease by Ultrastructure Defect and Genotype. Am J Respir Crit Care Med. 2019;199(2):190–8.

20. Ostrowski LE, Blackburn K, Radde KM, Moyer MB, Schlatzer DM, Moseley A, et al. A proteomic analysis of human cilia: identification of novel components. Mol Cell Proteomics. 2002;1(6):451–65.

21. Chivukula RR, Montoro DT, Leung HM, Yang J, Shamseldin HE, Taylor MS, et al. A human ciliopathy reveals essential functions for NEK10 in airway mucociliary clearance. Nat Med. 2020;26(2):244–51.

22. Blanchon S, Legendre M, Bottier M, Tamalet A, Montantin G, Collot N, et al. Deep phenotyping, including quantitative ciliary beating parameters, and extensive genotyping in primary ciliary dyskinesia. J Med Genet. 2020;57(4):237–44.

23. Brody SL, Pan J, Huang T, Xu J, Xu H, Koenitizer J, et al. Loss of an extensive ciliary connectome induces proteostasis and cell fate switching in a severe motile ciliopathy. bioRxiv. 2024.

24. Zawawi F, Shapiro AJ, Dell S, Wolter NE, Marchica CL, Knowles MR, et al. Otolaryngology Manifestations of Primary Ciliary Dyskinesia: A Multicenter Study. Otolaryngol Head Neck Surg. 2022;166(3):540–7.

25. Stoltz DA, Meyerholz DK, and Welsh MJ. Origins of cystic fibrosis lung disease. N Engl J Med. 2015;372(16):1574–5.

26. Rogers CS, Abraham WM, Brogden KA, Engelhardt JF, Fisher JT, McCray PB, Jr., et al. The porcine lung as a potential model for cystic fibrosis. Am J Physiol Lung Cell Mol Physiol. 2008;295(2):L240–63.

27. Rogers CS, Stoltz DA, Meyerholz DK, Ostedgaard LS, Rokhlina T, Taft PJ, et al. Disruption of the CFTR gene produces a model of cystic fibrosis in newborn pigs. Science. 2008;321(5897):1837-41.

28. Pezzulo AA, Tang XX, Hoegger MJ, Abou Alaiwa MH, Ramachandran S, Moninger TO, et al. Reduced airway surface pH impairs bacterial killing in the porcine cystic fibrosis lung. Nature. 2012;487(7405):109-13.

29. Tang XX, Ostedgaard LS, Hoegger MJ, Moninger TO, Karp PH, McMenimen JD, et al. Acidic pH increases airway surface liquid viscosity in cystic fibrosis. J Clin Invest. 2016;126(3):879–91.

30. Ostedgaard LS, Moninger TO, McMenimen JD, Sawin NM, Parker CP, Thornell IM, et al. Gel-forming mucins form distinct morphologic structures in airways. Proc Natl Acad Sci U S A. 2017;114(26):6842–7.

31. Hoegger MJ, Fischer AJ, McMenimen JD, Ostedgaard LS, Tucker AJ, Awadalla MA, et al. Impaired mucus detachment disrupts mucociliary transport in a piglet model of cystic fibrosis. Science. 2014;345(6198):818-22.

32. Stoltz DA, Meyerholz DK, Pezzulo AA, Ramachandran S, Rogan MP, Davis GJ, et al. Cystic fibrosis pigs develop lung disease and exhibit defective bacterial eradication at birth. Sci Transl Med. 2010;2(29):29ra31.

33. Zariwala MA, Leigh MW, Ceppa F, Kennedy MP, Noone PG, Carson JL, et al. Mutations of DNAI1 in primary ciliary dyskinesia: evidence of founder effect in a common mutation. Am J Respir Crit Care Med. 2006;174(8):858–66.

34. Zietkiewicz E, Nitka B, Voelkel K, Skrzypczak U, Bukowy Z, Rutkiewicz E, et al. Population specificity of the DNAI1 gene mutation spectrum in primary ciliary dyskinesia (PCD). Respir Res. 2010;11(1):174.

35. Guichard C, Harricane MC, Lafitte JJ, Godard P, Zaegel M, Tack V, et al. Axonemal dynein intermediate-chain gene (DNAI1) mutations result in situs inversus and primary ciliary dyskinesia (Kartagener syndrome). Am J Hum Genet. 2001;68(4):1030–5.

36. Wilkerson CG, King SM, Koutoulis A, Pazour GJ, and Witman GB. The 78,000 M(r) intermediate chain of Chlamydomonas outer arm dynein isa WD-repeat protein required for arm assembly. J Cell Biol. 1995;129(1):169–78.

37. Ostedgaard LS, Price MP, Whitworth KM, Abou Alaiwa MH, Fischer AJ, Warrier A, et al. Lack of airway submucosal glands impairs respiratory host defenses. Elife. 2020;9.

38. Labun K, Montague TG, Krause M, Torres Cleuren YN, Tjeldnes H, and Valen E. CHOPCHOP v3: expanding the CRISPR web toolbox beyond genome editing. Nucleic Acids Res. 2019;47(W1):W171–W4.

39. Whitworth KM, Benne JA, Spate LD, Murphy SL, Samuel MS, Murphy CN, et al. Zygote injection of CRISPR/Cas9 RNA successfully modifies the target gene without delaying blastocyst development or altering the sex ratio in pigs. Transgenic Res. 2017;26(1):97–107.

40. Yuan Y, Spate LD, Redel BK, Tian Y, Zhou J, Prather RS, et al. Quadrupling efficiency in production of genetically modified pigs through improved oocyte maturation. Proc Natl Acad Sci U S A. 2017;114(29):E5796–E804.

41. Chen PR, Redel BK, Spate LD, Ji T, Salazar SR, and Prather RS. Glutamine supplementation enhances development of in vitro-produced porcine embryos and increases leucine consumption from the medium. Biol Reprod. 2018;99(5):938–48.

42. Redel BK, Spate LD, and Prather RS. In Vitro Maturation, Fertilization, and Culture of Pig Oocytes and Embryos. Methods Mol Biol. 2019;2006:93-103.

43. Ye J, Coulouris G, Zaretskaya I, Cutcutache I, Rozen S, and Madden TL. Primer-BLAST: a tool to design target-specific primers for polymerase chain reaction. BMC Bioinformatics. 2012;13:134.

44. Shoemark A, Pinto AL, Patel MP, Daudvohra F, Hogg C, Mitchison HM, et al. PCD Detect: enhancing ciliary features through image averaging and classification. Am J Physiol Lung Cell Mol Physiol. 2020;319(6):L1048–L60.

45. Chen JH, Stoltz DA, Karp PH, Ernst SE, Pezzulo AA, Moninger TO, et al. Loss of anion transport without increased sodium absorption characterizes newborn porcine cystic fibrosis airway epithelia. Cell. 2010;143(6):911–23.

46. Jain A, Kim BR, Yu W, Moninger TO, Karp PH, Wagner BA, et al. Mitochondrial uncoupling proteins protect human airway epithelial ciliated cells from oxidative damage. Proc Natl Acad Sci U S A. 2024;121(10):e2318771121.

47. Hoegger MJ, Awadalla M, Namati E, Itani OA, Fischer AJ, Tucker AJ, et al. Assessing mucociliary transport of single particles in vivo shows variable speed and preference for the ventral trachea in newborn pigs. Proc Natl Acad Sci U S A. 2014;111(6):2355–60.

48. Bouzek DC, Abou Alaiwa MH, Adam RJ, Pezzulo AA, Reznikov LR, Cook DP, et al. Early Lung Disease Exhibits Bacteria-Dependent and -Independent Abnormalities in Cystic Fibrosis Pigs. Am J Respir Crit Care Med. 2021;204(6):692–702.

49. Adam RJ, Michalski AS, Bauer C, Abou Alaiwa MH, Gross TJ, Awadalla MS, et al. Air trapping and airflow obstruction in newborn cystic fibrosis piglets. Am J Respir Crit Care Med. 2013;188(12):1434–41.

50. Chang EH, Pezzulo AA, Meyerholz DK, Potash AE, Wallen TJ, Reznikov LR, et al. Sinus hypoplasia precedes sinus infection in a porcine model of cystic fibrosis. Laryngoscope. 2012;122(9):1898–905.

51. Chang EH, Tang XX, Shah VS, Launspach JL, Ernst SE, Hilkin B, et al. Medical reversal of chronic sinusitis in a cystic fibrosis patient with ivacaftor. Int Forum Allergy Rhinol. 2015;5(2):178–81.

52. Wee WB, Leigh MW, Davis SD, Rosenfeld M, Sullivan KM, Sawras MG, et al. Association of Neonatal Hospital Length of Stay with Lung Function in Primary Ciliary Dyskinesia. Ann Am Thorac Soc. 2022;19(11):1865–70.

53. Machogu E, and Gaston B. Respiratory Distress in the Newborn with Primary Ciliary Dyskinesia. Children (Basel). 2021;8(2).

54. Mullowney T, Manson D, Kim R, Stephens D, Shah V, and Dell S. Primary ciliary dyskinesia and neonatal respiratory distress. Pediatrics. 2014;134(6):1160–6.

55. Hyland RM, and Brody SL. Impact of Motile Ciliopathies on Human Development and Clinical Consequences in the Newborn. Cells. 2021;11(1).

56. Knowles MR, Daniels LA, Davis SD, Zariwala MA, and Leigh MW. Primary ciliary dyskinesia. Recent advances in diagnostics, genetics, and characterization of clinical disease. Am J Respir Crit Care Med. 2013;188(8):913–22.

57. Knowles MR, Zariwala M, and Leigh M. Primary Ciliary Dyskinesia. Clin Chest Med. 2016;37(3):449–61.

58. Bennett WD, Foster WM, and Chapman WF. Cough-enhanced mucus clearance in the normal lung. J Appl Physiol (1985). 1990;69(5):1670-5.

59. Sackner MA, and Kim CS. Phasic flow mechanisms of mucus clearance. Eur J Respir Dis Suppl. 1987;153:159–64.

60. Noone PG, Bennett WD, Regnis JA, Zeman KL, Carson JL, King M, et al. Effect of aerosolized uridine-5’-triphosphate on airway clearance with cough in patients with primary ciliary dyskinesia. Am J Respir Crit Care Med. 1999;160(1):144–9.

